# N-glycoproteomic profiling of the hookworm *Ancylostoma ceylanicum* reveals heterogeneity and phosphorylcholine-modified glycans in larval and adult parasites

**DOI:** 10.64898/2026.10.26.762015

**Authors:** Christina L. Paraggio, Vincent Chang, Keira E. Mahoney, Nara Chung, Sonali Sunsunwal, Stephanie Archer-Hartmann, Savanna Randi, Richard Bungiro, Lisa M. Harrison, Michel Ledizet, Parastoo Azadi, Michael Cappello, Stacy A. Malaker

## Abstract

Glycans of parasitic worms (helminths) have been previously described with roles in immunomodulation and the recognition of helminth proteins by the host immune system. Despite their significance in other systems, little is known about the glycosylation patterns of the hookworm *Ancylostoma ceylanicum*, an intestinal parasite of animals and humans, due to challenges in sample collection and structural complexity of many nematode glycans. Herein, we describe results of a released N-glycan analysis of adult hookworms, which revealed oligomannose, hybrid, and complex structures, including those with non-mammalian features. Next, we report the results of glycoproteomic analyses to identify hookworm glycoproteins and their associated glycan structures. We compared three enrichment methods and found that strong anion-exchange electrostatic repulsion liquid chromatography (SAX-ERLIC) led to the identification of the most N-glycopeptides. Analyses of larval and adult extracts revealed 82 larval and 157 adult N-glycoproteins, the majority of which were stage-specific. Of particular interest, phosphorylcholine (PC), a known immunomodulatory glycan modification, was detected in both stages, and several unique PC-modified structures were identified in each. Collectively, these results provide the first site-specific characterization of intact N-glycopeptides in *A. ceylanicum* and, with an adult-stage released glycan analysis, establish a method for glycoproteomic characterizations of other helminths.

## INTRODUCTION

Glycosylation, the addition of a sugar residue to a protein backbone, is one of the most common post-translational modifications that decorate eukaryotic proteins, Such modifications are generally N-linked, occurring at asparagine (N) residues of an N_X_S/T sequon (where X is any amino acid except proline), or O-linked, occurring on serine or threonine residues^1–3^. In addition to having integral roles in protein folding, stability, and function, the glycosylation patterns of proteins from eukaryotic parasites are important for development and host-pathogen interactions. Glycosylation of parasite proteins has been implicated in host Th2 immune priming, antibody recognition of parasitic worm (helminth) antigens, and altered cytokine responses^4–9^. In the case of *Haemonchus contortus*, an intestinal roundworm (nematode) parasite of sheep, native glycosylation patterns of the vaccine antigen H11 are critical to its ability to confer optimal immunoprotection^10,11^.

The hookworms *Ancylostoma ceylanicum, Ancylostoma duodenale*, and *Necator americanus* are, like *H. contortus*, Clade V nematodes. These are bloodfeeding helminths that mechanically attach to the intestinal epithelium of the host, causing iron deficiency anemia and growth delay in animal or human hosts^12–17^. Global burden of disease estimates from 2021 attribute 540,000 disability adjusted life years (DALYs) to hookworm infections, approximately 39.1% of DALYs from all intestinal nematode infections^18^. Hookworm infections are classified as a neglected tropical disease. Endemicity is associated with regions of rural poverty and underdeveloped sanitation practices^12,19^ as infected hosts release hookworm eggs in their feces. When these eggs hatch in the environment, larvae are released and molt to the infectious third stage (L3). The L3 penetrate the skin of a host, invade a blood vessel or lymphatic, and travel through the circulatory system to the pulmonary vasculature. After traversing the lung capillaries into the alveolar space, L3 migrate to the upper airway, where they are swallowed and enter the gastrointestinal tract. Of note, the larvae of *Ancylostoma* hookworms are also infectious by ingestion. During their migration, these larvae undergo a series of molts and ultimately reach the adult stage in the small intestine. Adult male and female hookworms mate in the intestine, leading to the release of eggs in host feces, restarting the life cycle^12^. In contrast to larvae, which exist in the environment and undergo multiple developmental molts, adult hookworms establish persistent infection within the mammalian host^12,20,21^. Given these differences, we set out to identify glycoproteomic signatures that mark changes in parasite development, with the longer-term goal of uncovering fundamental mechanisms of hookworm biology and pathogenesis.

Little is known about the glycosylation patterns of human hookworm proteins. Early studies revealed the presence of terminal sugars on the surfaces of the larval sheaths of hookworms by incubating larvae with carbohydrate binding proteins called lectins^22^. *In vitro* interactions between larvae of the hookworm *Necator americanus* with DC-SIGN and mannose receptor have suggested a role for this binding in larval exsheathment^23^. Genomic data from the hookworm *N. americanus* showed that adult worms express hexosyl transferases, a type of glycosyltransferase, at higher levels than larvae^24^, suggesting an increased role for protein glycosylation at later stages of hookworm development. Despite the fact that the proteomic landscape of *A. ceylanicum* has been recently investigated^25^, the glycome and glycoproteome remain uncharacterized, limiting our understanding of hookworm biology and pathogenesis. Putative N-glycosylation sites of excretory/secretory proteins of the related *Ancylostoma caninum* have been reported^26^, however this analysis was not intended to detect glycopeptides. Consequently, the specific glycan structures present in this species, as well as the site-specific localization of glycans on individual proteins, remain unknown. Progress in this area has been limited not only by the difficulty of obtaining sufficient parasite material across developmental stages, but also by the analytical complexity of nematode glycans, which display substantial structural diversity including features that are distinct from mammalian glycosylation patterns^27^. Evidence from other helminth systems suggests a role for parasite glycans in immunopathogenesis^5,6,8,28–35^, underscoring our interest in pursuing an in-depth characterization of hookworm glycosylation.

Herein, we describe the results of the first N-glycoproteomic profiling of the infectious (L3) and chronic (adult) stages of the hookworm *A. ceylanicum*. We first conducted a released glycan (glycomic) analysis of adult proteins, which revealed a variety of complex, hybrid, and mannosidic glycans, including those with non-mammalian features. Next, we screened larval and adult samples using an intact glycoproteomic workflow to gain insight into both the glycan structures and glycosylated proteins expressed at two key stages of parasite development. To capture hookworm glycopeptides for analysis, we compared 3 peptide-level enrichment methods: strong anion-exchange electrostatic repulsion liquid chromatography (SAX-ERLIC), PolyHYDROXYETHYL Aspartamide™ (PHEA), and iSPE® hydrophilic interaction liquid chromatography (Hilicon iSPE®). SAX-ERLIC performed the best, leading to the identification of the most glycopeptides and glycoproteins. We identified 82 N-glycoproteins in *A. ceylanicum* larvae, 157 in adult hookworms, and a total of 50 N-glycan structures using this workflow. These analyses suggest that microheterogeneity is a dominant feature of *A. ceylanicum* N-glycosylation both within and between stages. Phosphorylcholine, a known immunomodulatory glycan modification, was also identified on larval and adult proteins. Collectively, these findings highlight stage-divergent features of the *A. ceylanicum* N-glycoproteome. Future studies are needed to assess the roles of these and other glycans in hookworm immunopathogenesis.

## MATERIALS & METHODS

### *Ancylostoma ceylanicum* life cycle maintenance

Male weanling hamsters (21 days old) of the outbred *Mesocricetus auratus* HsdHan:AURA strain (Envigo, Indianapolis, IN) were infected with 100 third-stage *A. ceylanicum* larvae (L3) by oral gavage. Infection status was confirmed by enumerating the eggs per gram (EPG) of infected cages of hamsters using a modified McMaster method^28^. Fecal samples containing eggs were mixed with bone charcoal (Ebonex Corporation, Detroit, MI) and incubated for 1-2 weeks at 27°C. Third stage larvae (L3) were collected by Baermann funnel. At 21 days post-infection, hamsters were euthanized by CO_2_ inhalation, and adult worms were manually retrieved from the small intestines. During collection, the exteriors of the adult worms were stripped of all visible host material and rinsed with 1x phosphate buffered saline (PBS) to minimize contamination of downstream samples with host proteins.

### Preparation of hookworm extracts

We analyzed soluble hookworm proteins prepared from L3 stage larvae (<u>L</u>arval <u>Ex</u>tract, or LEX) or from pooled male and female adult worms (Adult <u>H</u>ookworm <u>Ex</u>tract, or HEX). *A. ceylanicum* larvae or adult worms were rinsed with 1x PBS and pulse-sonicated for 5 minutes at 40% amplitude. The sample was centrifuged at 14,000 rcf for 5 minutes to remove insoluble material, and the supernatant was concentrated approximately 10-fold^36–38^. The concentrations of each protein sample were determined by Pierce BCA assay (ThermoFisher, Waltham MA). All samples were stored at –80 °C prior to use.

### Release of N-glycans from adult hookworm extract (HEX)

Following chloroform:methanol precipitation, 1 mg of adult *A. ceylanicum* protein extract (HEX) dissolved in 50 mM ammonium bicarbonate, was exchanged with the same buffer to eliminate residual solvents that might interfere with glycan release and downstream analysis. Protein concentration was quantified by BCA assay, and 400 μg of protein was used for subsequent analysis. The proteins were reduced using 5 mM DTT (dithiothreitol) and incubated at 45 °C for 45 min. Next, the proteins were alkylated using 10 mM IAA (iodoacetamide) and incubated for 20 min at room temperature. DTT and IAA were removed by desalting against 50 mM ammonium bicarbonate (NH_4_HCO_3_) with 3-kDa cutoff centrifugal filters. After desalting, 3 μL of PNGase F and 3 μL of PNGase A were added to the sample and incubated at 37 °C for 48 h. After incubation, the released N-glycans were recovered by passing the sample through 10-kDa cutoff centrifugal filters and collecting the flowthrough. The filtrate (containing N-glycans) was split in half and lyophilized. One fraction was processed for permethylation^39^ and the other for procainamide labeling.

### Permethylation of released N-glycans from HEX

N-glycans were permethylated as previously described^40^. First, 100 μL of NaOH was added to prewashed glass tubes, followed by 200 μL of MeOH. Tubes were vortexed, and 4 mL of DMSO was added prior to subsequent vortexing. Tubes were then centrifuged at ∼1100 rcf, and the upper liquid layer removed, keeping the gel at the bottom intact. This procedure was repeated 5 times before adding 1 mL of DMSO. Lyophilized N-glycans were dissolved in 200 μL of DMSO followed by 300 μL of freshly prepared DMSO/NaOH base and 100 μL of iodomethane. The resulting mixture was sonicated for 15 minutes. Another 300 μL of base was added, together with 100 μL of iodomethane, prior to additional sonication (15 minutes). Water (2 mL) was added to quench the reaction, followed by 3 mL of dichloromethane (DCM) to extract N-glycans with vigorous shaking. The resulting sample was centrifuged for 1 min and the upper fraction removed. This procedure was repeated 4 more times. The lower DCM layer was then transferred to a fresh tube and dried under nitrogen gas.

### Procainamide labeling of released N-glycans from HEX

Fresh stocks of procainamide (55 mg in 100 μL DMSO) and sodium cyanoborohydride (22 mg in 100 μL water) were prepared. The procainamide solution was acidified by mixing 1-part glacial acetic acid with 8 parts stock solution. To the lyophilized N-glycans, 18 μL of prepared acidified procainamide label and 24 μL of the freshly prepared cyanoborohydride reagent were added, mixed well and incubated at 37 °C for 16-20 h. After incubation, the sample was diluted with water in the ratio 1:10, filtered, and 2 μL of the diluted sample was injected on an Orbitrap Ascend Tribrid mass spectrometer.

### Glycomic data acquisition from HEX

Dried, permethylated glycans were reconstituted in 20 μL of MeOH. Each sample was mixed with 2,5-dihydroxybenzoic acid (DHB) matrix in a 1:1 ratio and ∼0.6 μL of the mixture was spotted on MTP 384 target plate polished steel. After drying up the spots, MALDI-MS1 data was acquired on a Bruker Rapiflex MALDI-TOF. For determining glycan structures, including modifications that would be impacted by permethylation, LC-MS/MS on the procainamide labeled N-glycans was carried out. MS2 data from the procainamide labeled N-glycans was manually assessed with assistance from Glycoworkbench v2.0.

### Preparation and purification of tryptic peptides for comparison of enrichment methods

Proteins from LEX (0.826 μg/μL) was purified by chloroform:methanol precipitation. First, 400 μL of MeOH was added to 100 μL of extract. The sample was then vortexed and incubated at –20°C for 1 h. Next, 100 μL of chloroform was added, then 300 μL of LC-MS grade H_2_O with vortexing between each addition. The samples were centrifuged at 14,000 rcf for 1 min, and the aqueous layer was discarded. To the proteinaceous sample, 400 μL of MeOH was added. The sample was vortexed and centrifuged at 20,000 rcf for 5 minutes. Methanol was removed and the protein pellet was air-dried prior to storage at –80°C. The dried sample was reconstituted with 100 μL of 20 mM Tris (pH 8). The sample was reduced by adding 4 μL of 25 mM DTT and incubating at 65 °C for 20 min on a shaker at 1200 rpm (Eppendorf, ThermoMixer C). After cooling to room temperature, 8 μL of 35 mM IAA was added. The sample was incubated at room temperature in the dark for 15 min, then stored at –20 °C. The sample was digested with trypsin (0.05 μg/μL) at a ratio of 1:200 and was incubated at 37 °C overnight. The next morning, the reaction was quenched with 1.5 μL of formic acid. Samples were desalted using 10 mg Strata-X 33 μm polymeric reversed phase SPE columns (Phenomenex, 8B-S100-AAK). Next, 200μL of 0.5% formic acid was added to each sample and vortexed. Columns were primed with 1 mL of acetonitrile (ACN), followed by 1 mL of 0.1% formic acid in H_2_O, and then by 1 mL of 40% ACN with 0.1% formic acid in H_2_O. The sample was transferred to the column. Two 150 μL volumes of 0.1% formic acid were used to rinse the sample tube and were subsequently loaded onto the column. Samples were eluted with two 150 μL additions of 40% ACN 0.1% formic acid, and the eluates were vacuum-dried. Samples were reconstituted with 100 μL of 0.1% formic acid and split into 3 aliquots of equal volume for use in each enrichment method as follows.

### Strong anion-exchange electrostatic repulsion liquid chromatography (SAX-ERLIC) and iSPE® hydrophilic interaction liquid chromatography (Hilicon iSPE®)

Thermo HyperSep™ Retain AX cartridges (Thermo Scientific, 60107-403) or iSPE®-HILIC (Hilicon iSPE®) columns were activated with 2 mL of LC-MS grade H_2_O, followed by 1 mL of 100 mM triethylammonium acetate, and two volumes of 1 mL 95% ACN with 0.1% trifluoroacetic acid (TFA). Columns were conditioned with two volumes of 1 mL of 95% ACN with 1% TFA. Two aliquots of ∼30 μg LEX prepared as described, one for each column, were brought up in 1 mL of 95% ACN with 1% TFA and loaded onto the columns. Sample tubes were rinsed with 1 mL of the same and loaded onto the columns. Glycopeptides were eluted with two additions of 500 μL of 40% ACN with 0.1% TFA. Samples were dried and stored at –20° C.

### PolyHYDROXYETHYL Aspartamide™ (PHEA) glycopeptide enrichment

Unfiltered pipette tips (200 μL) were plugged with Whatman filter membranes. PolyHYDROXYETHYL Aspartamide™ (PHEA) (PolyLC®, BMHY12-03) packing material (12 μm, 300 Å) was added to a tube. A packing slurry was created by adding 500 μL of 200 mM ammonium formate (AF), 100 μL of which was added to each prepared pipette tip. Tips were spun for at 350 rcf for approximately 2 mins until the solution was almost gone, leaving 5-10 mm of packed PHEA material. Between each step, tips were centrifuged at 350 rcf until the mobile phase flowed through almost entirely. Tips were rinsed as follows: 100 μL of 1% formic acid, 100 μL of 20 mM AF, 90% ACN, 100 μL of 200 mM AF, 100 μL of H_2_O, then 2x with 100 μL of 20 mM AF, 90% ACN. One aliquot of desalted LEX was vacuum-dried to remove the 0.1% formic acid and was reconstituted by first adding 10 μL of 200 mM AF, vortexing, then adding 90 μL of ACN. Samples were added to the prepared tips, which were rinsed 2x with 100μL of 20 mM AF, 90% ACN to elute small molecules and salts. Unmodified peptides were eluted with two additions of 100 μL 40 mM AF, 80% ACN. These samples were collected, dried down, and stored at –80°C. Glycopeptides were eluted as follows. First, 150 μL of 1% formic acid was added and the sample centrifuged at 200 rcf for 5 minutes. A second volume of 100 μL of 1% formic acid was added and centrifuged at 200 rcf until the mobile phase flowed through completely. Enriched samples were dried and stored at –20 °C.

### Identification of N-glycopeptides by LC-MS/MS

Samples were analyzed by online nanoflow liquid chromatography-tandem mass spectrometry using an Orbitrap Eclipse Tribrid mass spectrometer (Thermo Fisher Scientific) coupled to a Dionex UltiMate 3000 HPLC instrument (Thermo Fisher Scientific). Samples were resuspended in 8 μL of buffer A (0.1% formic acid), and 6 μL were injected onto an Acclaim PepMap 100 column packed with 2 cm of 5 μm C18 material (Thermo Fisher, 164564). Peptides were then separated on a 15 cm PepMap RSLC EASY-Spray C18 column packed with 2 μm C18 material (Thermo Fisher, ES904) using a gradient from 0 to 35% solvent B (0.1% formic acid with 80% ACN) in 60 min with a constant flow rate of 300 nL min-1. Full scan MS1 spectra were acquired at a resolution of 60,000 full width at half maximum (FWHM) at 400 m/z, an automatic gain control (AGC) target of 3e5, and a mass range from 300 to 1,500 m/z. Dynamic exclusion settings allowed 2 repeats in 7 s before exclusion for 7 s. Only charge states 2 to 6 were selected for fragmentation. MS2s were generated at top speed for 3 s. Fragmentation scans were collected in the Orbitrap at 15,000 resolution. Higher-energy collisional dissociation (HCD) was performed on all selected precursor masses with the following parameters: isolation window of 2 m/z, stepped collision energy 25 – 30–40% normalized collision energy (nCE), Orbitrap detection (resolution of 7,500), maximum inject time of 50 ms, and a standard AGC target. An additional electron transfer dissociation (ETD) fragmentation of the same precursor was triggered if 1) 3 of 8 HexNAc or NeuAc fingerprint ions (126.055, 138.055, 144.07, 168.065, 186.076, 204.087, 274.092, and 292.103) were present at ± 0.1 m/z and greater than 5% relative intensity, 2) the precursors were present at a sufficient abundance for localization using ETD. If the precursor was between 300–850 m/z and over 3e4 intensity, an ETD scan was triggered and read out using the ion trap at a normal scan rate. For precursors between 850–1500 m/z with an intensity greater than 1e5, supplemental activation was applied at 15% nCE and read out in the Orbitrap at 7500 resolution. Both used charge-calibrated ETD reaction times, 150 ms maximum injection time, and 200% standard injection targets. The mass spectrometry proteomics data have been deposited to the ProteomeXchange Consortium via the PRIDE^41^ partner repository with the dataset identifier PXD085231.

### Glycoproteomic data analysis

Due to the dearth of available data on the glycosylation patterns of human hookworms, and to account for nematode-specific glycan motifs, an N-glycan library was compiled from published reports characterizing the following helminth species: *Fasciola hepatica*^42^*, Dirofilaria immitis*^28^*, Haemonchus contortus*^43^, and *Schistosoma mansoni*^30^. Glycans identified from the Clade V nematode *Caenorhabditis elegans* ^44^ were also included, although it is not a helminth. The Byonic software package was used to search raw files against the *A. ceylanicum* proteome (UniProt) to generate a focused protein library. This library was used with the in-house compiled N-glycan database to search the raw data with the following considerations. Carbamidomethyl Cys was set as a fixed modification, and results were searched using ‘fully specific’ C-terminal cleavage to Lys or Arg with a maximum of 2 missed cleavages allowed. One rare modification and a maximum of 2 common modifications were allowed. Search results were filtered with a Byonic score of >150 and LogProb >1.5. Results were sorted by peptides with an identified glycan present, and duplicates were removed to identify unique glycopeptides. To assess potential host contamination, one sample of SAX-ERLIC enriched adult hookworm proteins was additionally searched against a combined proteome of reviewed *M. auratus* proteins (UniProt) and the focused *A. ceylanicum* library. Glycans were omitted from this search, and decoys and contaminates were included. Manual validation of selected structures was conducted by identifying diagnostic fragment ions in the MS2 output for these glycopeptides.

### Final workflow for glycoproteomic sample preparation

Following the comparison of enrichment methods, 3 replicates of 100 μg *A. ceylanicum* LEX and HEX proteins were prepared by reduction and alkylation, chloroform:methanol precipitation, and desalting, and were then vacuum-dried. SAX-ERLIC was performed using a modified protocol as follows. Samples were reconstituted in 10 μL of 0.1% TFA in LC-MS grade H_2_O, vortexed and spun down. One third of this sample was taken for enrichment, to which 1 mL of 95% ACN in 1% aqueous TFA was added. Thermo HyperSep™ Retain AX cartridges (Thermo Scientific, 60107-403) were activated with 3 mL of LC-MS grade MeOH, followed by 3 mL of H_2_O, and 3 mL of triethylammonium acetate buffer (Sigma 90358) diluted 1:10 in LC-MS grade H_2_O. Columns were equilibrated with 3 mL of 95% ACN in 1% aqueous TFA, followed by 3 mL of 40% ACN in 0.1% aqueous TFA, and 3 mL of 95% ACN in 1% aqueous TFA. Samples were loaded onto the columns, and the columns were washed with 1 mL of 95% ACN in 1% aqueous TFA. Enriched glycopeptides were eluted with 2 volumes of 500 μL of 40% ACN in 0.1% aqueous TFA and vacuum dried. Samples were dried in a vacuum and resuspended in 8 μL of 0.1% formic acid, 6 μL of which were injected onto the Orbitrap Eclipse Tribrid mass spectrometer (Thermo Fisher Scientific). LC-MS/MS and searching were performed as described previously. The final list of glycopeptides and glycoproteins represents those identified in 2 out of 3 replicates. Data was visualized by GraphPad PRISM (v 11.1.0) to generate plots and the *eulerr* package in R (v 4.3.1) to generate Euler diagrams.

### MSFragger and FragPipe analyses

MSFragger was used to generate a Pearson correlation comparing larval (LEX) and adult (HEX) pooled samples. Raw data files were uploaded as two biological samples with 3 technical replicates each, along with the *A. ceylanicum* UniProt library, and compiled N-glycan library. Analysis was conducted using the FragPipe computational platform to obtain peptide identifications and their respective label-free quantification values using IonQuant. Statistical analysis was performed based on the combined peptide.tsv file. First, contaminants were filtered out. In addition, peptides that were not identified/quantified consistently in the same condition were removed. The quantification values were converted to log2 scale if intensities were used. A normalization was then applied. Samples were grouped by conditions, and missing values were imputed. Peptide-wise linear models combined with empirical Bayes statistics were used for the differential expression analyses. The limma package from R Bioconductor was used to generate a list of differentially expressed peptides for each pair-wise comparison. A cutoff of the adjusted p-value of 0.05 along with a |log2 fold change| of 1 was applied to determine differentially expressed peptides in each pairwise comparison. The parameters used are as follows: Normalization = none, Imputation = man, Tested comparisons = HEX_vs_LEX, Adjusted p-value cutoff <= 0.05, Log2 fold change cutoff >= 1.

## RESULTS

### Glycomic analysis of adult hookworms reveals structurally diverse N-glycans

A released N-glycan analysis was performed on soluble adult hookworm homogenate (HEX) to gain structural insight into the glycans expressed by mature *A. ceylanicum* worms. Permethylation and procainamide labeling were used in parallel to capture glycans across a range of abundances, and these experiments allowed us to identify complex, hybrid, paucimannose, and oligomannose structures (Figure 1). Procainamide labeling allowed for the detection of several additional species, among them structures with labile modifications that were not retained through permethylation. All glycans identified by either approach are summarized in Supplemental Table 1. The MALDI spectrum depicted in Figure 1 was dominated by ions corresponding to paucimannose glycans, with GlcNAc_2_Man_3_ (m/z 1171.517) giving the most intense signal. Ions for paucimannose structures (GlcNAc_2_Man_3–4_ ± core modifications) were prominent across the low-mass region, whereas complex and hybrid species were detected as comparatively weak signals at higher m/z. Relative peak intensities in permethylation MALDI are influenced by ionization efficiency and are therefore not strictly quantitative, so these signals were interpreted as an indication of which structures were readily detected rather than of their relative amounts. Two features not found in mammals were also apparent. Galactosylated fucose was detected on 3 structures (m/z 1345.573, 1549.636 and 2039.827) (Figure 1, inset), and phosphorylcholine (PC) was detected on 7 structures by procainamide labeling (Supplemental Table 1). Core modifications such as fucosylation have been previously reported in other helminths, some with immunomodulatory roles^27,45–50^. The detection of a single sialylated glycan (m/z 2605.151) may be a contaminating host glycan originating from the bloodmeal of adult worms, as sialic acid residues are present on erythrocytes^51^. This glycan was found by permethylation analysis alone and does not have MS2 data, thus other structures are also possible.

**Figure 1:**
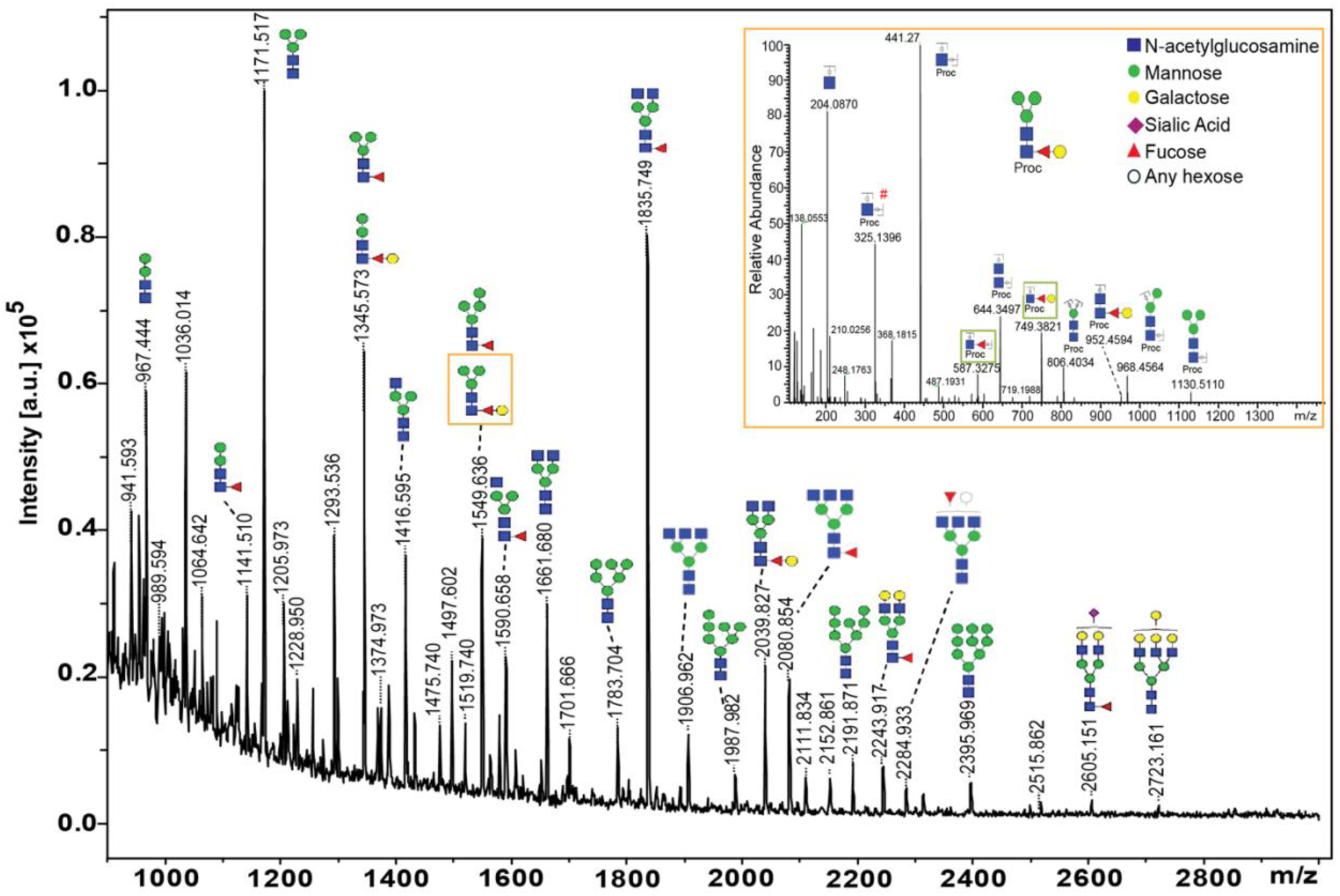
Released glycan analysis of adult proteins reveals diverse structural classes of N-glycans. MALDI-MS spectrum of highly abundant permethylated N-glycans from adult *A. ceylanicum*. All peaks are N-glycan plus Na adduct: [M+Na]^+^. Paucimannose (GlcNAc_2_Man_3–4_ ± core modifications) glycans were abundant in the low-mass region, whereas ions for hybrid and complex structures were identified at higher m/z. Modifications of the chitobiose core structure (GlcNAc_2_Man_3_) with fucose (red triangle) and galactosylated fucose (red triangle, yellow circle) were identified across the spectrum (m/z: 1141.510, 1345.573, 1549.636, 1590.658, 1835.749, 2039.827, 2080.854, 2243.917, and 2605.131). Galactosylated fucose structures are non-mammalian signatures and were additionally validated by procainamide-labeled MS2 data (orange box, inset) by the fragment ions m/z 587.3275 and 749.3821. Five structures (m/z: 1519.740, 1906.962, 2243.917, 2605.151, and 2723.161) were identified through permethylation alone.

### SAX-ERLIC enrichment leads to the greatest increase in glycosylation signal from larval proteins

Following the glycomic identification of non-mammalian N-glycans in adult *A. ceylanicum* homogenate, we set out to optimize an intact glycoproteomic workflow for use with hookworm proteins. By assessing the relative glycosylation signals, total glycopeptide identifications, and breadth of glycan structures, we first compared the ability of three glycopeptide enrichment methods (SAX-ERLIC, PHEA, Hilicon iSPE®) to retain larval hookworm (LEX) glycopeptides for analysis. These methods were selected because they are less biased towards specific glycan structures compared to lectin-based enrichment strategies^52^. An additional sample of LEX was prepared following the same workflow, but with the enrichment steps omitted to act as a control. To assess differences between enrichment methods, we compared the relative abundances of the oxonium marker ion for glycosylation (HexNAc^+^, m/z: 204.0867) from each sample. The overlay shown in Figure 2A suggests that many of the same peaks were retained between methods, except for peaks detected in the sample enriched by SAX-ERLIC (Figure 2A, coral) between 32.27-38.22 minutes. From this scan time range, 49 glycopeptides were identified in the SAX-ERLIC sample, compared to 15 in the Hilicon iSPE® sample, and 14 in the PHEA sample, indicating that these peaks corresponded to identified glycosylated species. However, peak intensity from 55.31-56.63 minutes appeared much greater in the samples enriched by PHEA (Figure 2A, pale blue) and Hilicon iSPE®) (Figure 2A, light green) compared to SAX-ERLIC (Figure 2A, coral), suggesting differences in the retention of certain analytes.

**Figure 2:**
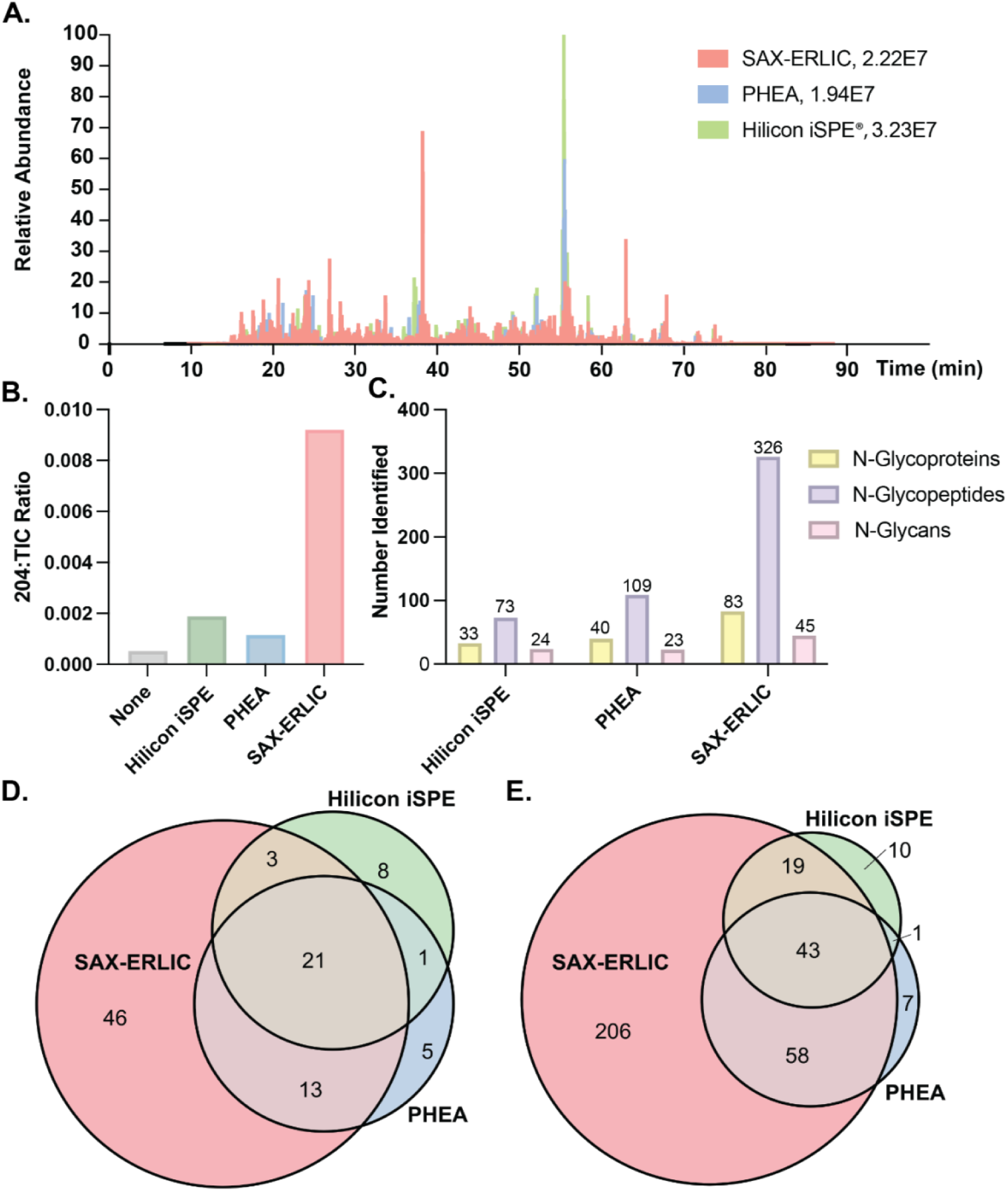
SAX-ERLIC outperforms other HILIC enrichments for the detection of glycosylation in *A. ceylanicum* larval proteins. **a**) Overlay of normalized extracted ion chromatograms (XIC) for HexNAc^+^ 204.0867 m/z from Xcalibur software. Similarities between peak distribution suggest retention of common analytes, with SAX-ERLIC (light red) leading to the detection of the majority of species. (b) Ratios of 204.0867 m/z marker ion for glycosylation compared to Total Ion Current (TIC) for each glycopeptide enrichment method tested, and one unenriched sample (“none”, gray). Ratios (204.0867: TIC) for each sample are as follows: None, 1.56E6: 2.96E9 (gray); Hilicon iSPE®, 3.23E7: 1.71E10 (light green); PHEA, 1.94E7; 1.69E10 (pale blue); SAX-ERLIC, 2.22E7: 2.41E9 (coral). (c) Numbers of unique larval N-glycoproteins, N-glycopeptides, and N-glycans identified in each enrichment method (Score >150, LogProb >1.5). Euler diagrams depicting overlap in (d) N-glycoproteins and (e) N-glycopeptides identified in each enrichment method. In both cases, SAX-ERLIC (coral) led to the identification of the majority of all detected glycosylated species, in addition to glycoproteins and glycopeptides that were not retained for analysis by other methods.

Enrichment with SAX-ERLIC also led to the highest relative glycosylation signal, determined by the ratio of the 204.0867 m/z signal to the total ion current (TIC), (Figure 2B, coral), Compared to the sample of unenriched pooled LEX, this constituted a 15.88-fold increase in relative glycosylation signal. Enrichment by PHEA and Hilicon iSPE® led to 1.98– and 3.26-fold increases, respectively (Figure 2B, Supplemental Table 2). This result reflects a more effective depletion of unmodified peptides by SAX-ERLIC. To determine the extent to which differences in the extracted ion chromatograms from each enrichment affected the identification of N-glycosylated species, we next compared the glycopeptides identified from each. Across all methods, 344 unique glycopeptides (UGPs) were identified (Supplemental Table 3). The vast majority (326) were identified following SAX-ERLIC enrichment (Figure 2C, lilac). PHEA and Hilicon iSPE®) led to identification of fewer glycopeptides (109 and 73, respectively) (Figure 2C, lilac). As expected, a similar trend was observed when comparing the number of identified N-glycoproteins (Figure 2C, yellow). These glycoproteins were largely representative of all those identified across the different enrichment methods (Figure 2D, red), with only 14 additional glycoproteins identified by PHEA (Figure 2D, blue) and Hilicon iSPE® (Figure 2D, green) combined. A comparison of the glycopeptides revealed similar results: 94.8% of those identified were included in the SAX-ERLIC enriched sample (Figure 2E, red). Lists of the identified glycoproteins by each method can be found in Supplemental Table 4. As observed with N-glycopeptides and N-glycoproteins, 93.75% (45/48) of identified N-glycans were found in the sample enriched by SAX-ERLIC (Figure 2C, pink, Supplemental Table 5). Although each enrichment was performed once on a single aliquot of pooled larval extract and these comparisons are qualitative, we found these results sufficiently convincing to move forward with SAX-ERLIC for subsequent glycoproteomic analyses.

### Identified N-glycoproteins are largely stage-specific in *A. ceylanicum* hookworms

A total of 320 glycopeptides were identified in larvae (82 N-glycoproteins) and 426 in adults (157 N-glycoproteins) (Figure 3A). These numbers represent those identified in 2 out of 3 replicates. The values for each individual sample are shown in Figure 3B. The identification of stage-specific hookworm proteins may have downstream applications as diagnostic markers of infection, especially if decorated with non-mammalian glycans that could differentiate a parasite from its mammalian host. Just 32 N-glycoproteins were common between stages, with 50 glycoproteins unique to larvae, and 125 unique to adults (Figure 3C, Supplemental Table 6). A comparison of glycopeptides showed a similar trend (Figure 3D, Supplemental Table 7). However, because more glycopeptides were identified in adults overall, we cannot exclude that some apparent larval-specific absences reflect detection depth rather than expression. A Pearson correlation of log2 intensities further confirms that, while replicates for the same sample type were highly similar (Figure 3E, top left, bottom right), adult and larval samples were distinct (Figure 3E, top right, bottom left). These findings suggest an element of stage specificity to the expression of hookworm glycoproteins. Stage-specific expression of glycoproteins appears to be a common feature of nematode development^30,31,44,53^ and likely reflects the different niches occupied throughout parasite lifecycles. Due to the physical attachment of adult hookworms to the intestinal epithelium and ingestion of host blood, we cross-referenced one replicate of adult proteins against a combined host and parasite proteome, and no glycoproteins were identified (Supplemental Table 8). To further assess the extent of stage-specific protein expression, we additionally compared the unenriched samples of larval and adult proteins. As expected, this analysis yielded significantly more proteins than our glycoproteomic workflow: 966 in larvae and 1442 in adults (Supplemental Table 9). Just 55 peptides (out of 6141) in larvae and 58 (out of 9043) in adults were identified with glycan structures (Supplemental Table 10). This further emphasizes the utility of the workflow reported above, and the necessity of including a peptide-level enrichment for glycoproteomic analysis. The relative proportion of common proteins increases in the proteome compared to the glycoproteome at both stages, from 39% (32/82) to 53.8% (520/966) of larval proteins and from 20.4% (32/157) to 36.1% (520/1442) of adult proteins (Supplemental Table 9). Together, these findings suggest that the hookworm N-glycoproteome is more stage-divergent than the proteome itself.

**Figure 3:**
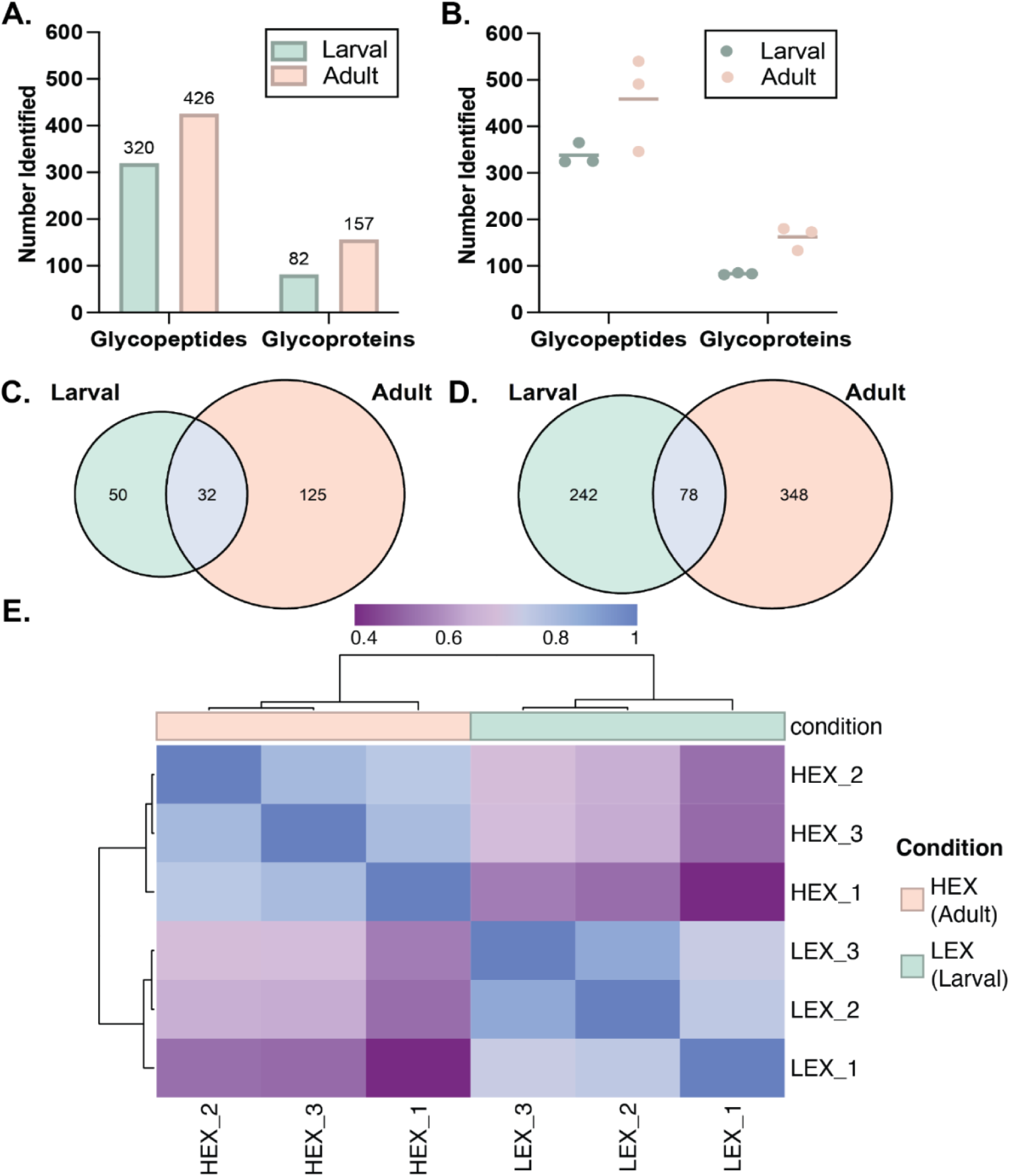
N-glycoproteins from larval and adult hookworms cluster by developmental stage. Three technical replicates of pooled larval (LEX) and adult (HEX) *A. ceylanicum* proteins were processed using the final glycoproteomic workflow described in Methods. To generate a final list of glycopeptides and glycoproteins, the data from each sample was sorted by Byonic Score >150 and LogProb >1.5. The resulting data were filtered to include only peptides with glycans, and duplicates were removed by starting position and calculated mass. The list of glycopeptides from each replicate was compared and a final list of glycopeptides was generated based on those found in 2 out of 3 replicates. This list was used in turn to determine the final list of glycoproteins and N-glycan structures. (a) Final numbers of N-glycoproteins and N-glycopeptides for both larval (mint) and adult (peach) *A. ceylanicum* proteins. (b) N-glycoproteins and N-glycopeptides identified in each replicate of larval (mint) and adult (peach) proteins. Final identifications were compared between stages, as represented in the Euler diagrams of (c) N-glycoproteins and (d) N-glycopeptides. A Pearson correlation was generated in FragPipe and MSFragger comparing all replicates to each other (e). Replicates within one sample type are highly similar, but those between sample types are distinct, concordant with our prior analyses.

### Stage-specific N-glycans decorate larval and adult hookworm proteins

The N-glycans decorating hookworm glycoproteins spanned mannosidic (oligomannose, paucimannose), complex, and hybrid structures. A total of 50 unique structures were identified: 35 in larval and 42 in adult parasites (Figure 4A, Supplemental Table 11). As observed with glycopeptides and glycoproteins (Figure 3B), the results between replicates were consistent for each sample type (Figure 4B). That several of the 32 common glycoproteins were predicted glycosidases and glycosyl hydrolases (Supplemental Table 6) suggested shared glycosylation machinery throughout parasite development. Indeed, comparing the glycan structures revealed that 54% (27/50) were identified in both larval and adult samples (Figure 4C). The remaining structures were identified in only one developmental stage. The 8 glycans that were identified in larvae alone were hybrid, complex, and paucimannose. Interestingly, 3/8 (37.5%) of these glycans were methylated. In other parasitic infections, methylated glycans are recognized by host antibodies and lectins^54,55^. A single methylated N-glycan structure was detected in the adult sample but was not unique to this stage. The potential of such glycans to serve as early-stage marker of hookworm infection remains to be studied. In both larvae and adults, oligomannose glycans, including paucimannose, were the most common, followed by complex, then hybrid (Figure 4D). These similar distributions indicate no preference for the major classes of N-glycans between larval and adult parasites. Among those oligomannose glycans, the majority (larvae: 14/19, adult: 11/17) were paucimannose (HexNAc_2_Man_3-4_ ± core modifications). These findings support our glycomics data, which showed that *A. ceylanicum* adults synthesized truncated, paucimannose N-glycans (Figure 1, Supplemental Table 1). In adults, 10/15 (66.67%) of the unique glycans were complex, which may suggest an increased role for such structures in the chronic stage of infection. Classifications of identified N-glycans can be found in Supplemental Table 12. Due to the electrostatic repulsion component of the SAX-ERLIC method, and the identification of a sialylated glycan structure by glycomics, we anticipated detecting a greater number of negatively charged or sialylated glycan structures by this method. However, none were identified in this or any of the enrichments, despite their inclusion in our compiled glycan database (Supplemental Figure 1). These results support previous reports that helminths lack pathways to synthesize sialic acid *de novo*^29,56^ and suggest an alternate structure for the single sialylated glycan proposed by permethylation analysis (Figure 1, m/z 2605.151).

**Figure 4:**
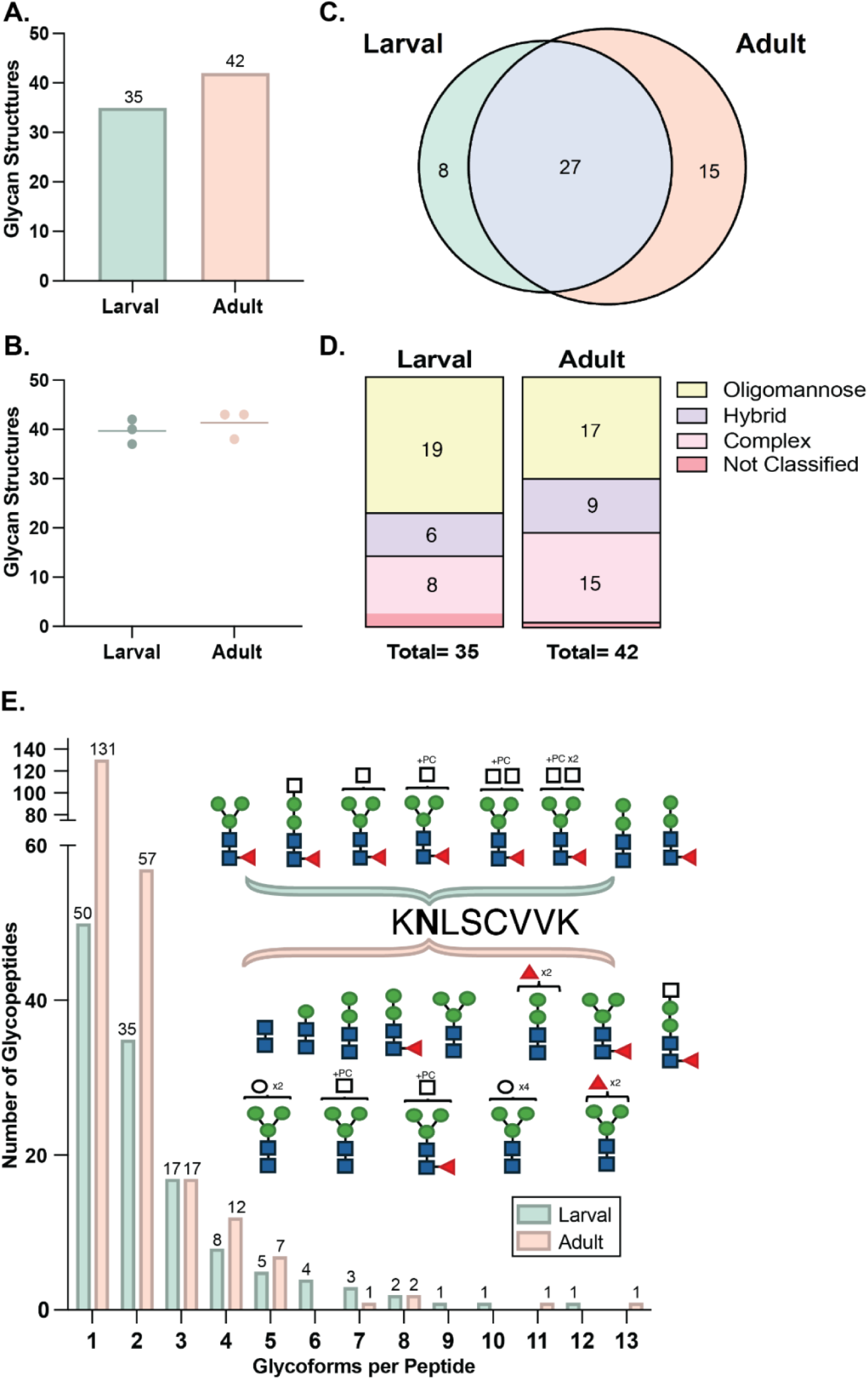
N-glycans identified by glycoproteomic workflow span mannosidic, hybrid, and complex structures. Glycan compositions were extracted from the lists of final glycopeptides at each stage. Duplicates were removed to generate lists of larval and adult glycans, numbers for which are shown in (a). Numbers of unique structures identified in each of the triplicate samples are shown in (b). A comparison of the final structures identified between stages is shown by Euler diagram in (c) and suggests both conserved glycosylation machinery and stage-specific structures. N-glycans are classified as oligomannose, complex, or hybrid. Oligomannose glycans have branches extended by mannose only, whereas the branches of complex glycans are comprised of other hexose sugars. Hybrid glycans have one branch comprised of mannose residues, and one branch extended by other sugars. Searched data from larval and adult confirmed glycan composition (HexNAc_X_Hex_Y_) but not structural data. As such, predicted structures based on these compositions were used to classify N-glycans into the major categories (d). N-glycans at both stages were predominantly oligomannose glycans, including paucimannose structures, although complex and hybrid glycans were identified in each sample as well. Differences in glycan structure at a given N_X_S/T sequon, or microheterogeneity, was a common feature in larval and adult hookworm proteins. Peptide backbones were extracted from the final lists of glycopeptides at each stage, and the numbers of glycoforms counted. The distribution of glycoforms per peptide backbone is shown in (e). The inset depicts an example of glycoforms identified for one such peptide, (K**N**LSCVVK) from the FYVE-type domain-containing protein A0A016UL41. Glycan structures were predicted by composition and the chitobiose core structure following the symbolic lexicon described in Figure 1. The determination of core fucose modifications is based on reports of similar structures in nematodes and their detection in glycomics data of adult hookworm proteins (Figure 1).

### Hookworm glycoproteins exhibit heterogeneous N-glycosylation

Changes in protein glycosylation throughout parasite development has been reported in several species^44,57–59^. We first compared the glycosylation patterns of the 32 glycoproteins identified in both larval and adult hookworms. Microheterogeneity, that is, variability in the glycan structures occupying the same N_X_S/T sequon of a peptide, was a salient feature. A total of 166 larval and 126 adult glycopeptides were identified corresponding to these common proteins, and just 69 were equivalent between stages (peptide + glycan) (Supplemental Table 13). Notably, all glycoforms with complex N-glycans were stage-specific (Supplemental Table 12). This may suggest a role for complex glycan structures in refining protein interactions throughout parasite development and merits future study. We next assessed macroheterogeneity, i.e., whether the same glycosites were occupied in larvae and adults. In 7 of the 32 common proteins (21.9%), the total number of occupied glycosites differed between stages (Table 1). Changes in the glycosylation patterns of these proteins may indicate a role for glycans in the fine-tuning of host-parasite interactions beyond protein expression.

**Table 1:**
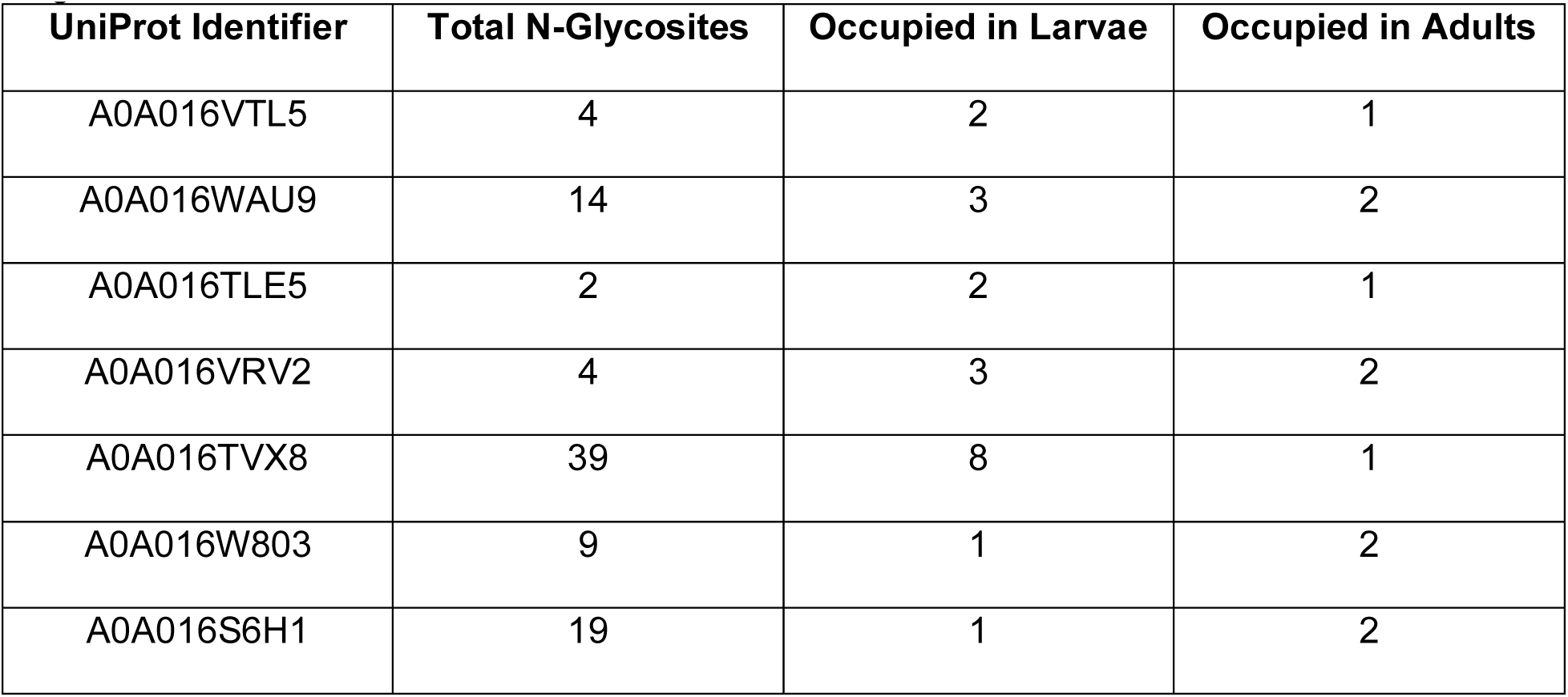
Macroheterogeneity in N-glycosylation is observed between hookworm developmental stages.

| UniProt Identifier | Total N-Glycosites | Occupied in Larvae | Occupied in Adults |
| --- | --- | --- | --- |
| A0A016VTL5 | 4 | 2 | 1 |
| A0A016WAU9 | 14 | 3 | 2 |
| A0A016TLE5 | 2 | 2 | 1 |
| A0A016VRV2 | 4 | 3 | 2 |
| A0A016TVX8 | 39 | 8 | 1 |
| A0A016W803 | 9 | 1 | 2 |
| A0A016S6H1 | 19 | 1 | 2 |

As heterogeneity was a feature of those proteins common to larval and adult hookworms, we next expanded our analysis to include the glycoproteins that were unique to either stage. We found 127 unique peptide backbones in the larval sample (320 total glycopeptides) and 229 in adults (426 total glycopeptides). While most backbones had 1-3 glycoforms, there were a subset of peptides at each stage with ≥10 glycoforms (Figure 4E, Supplemental Tables 14, 15). The peptides with the maximum number of glycoforms per stage were different between samples. In adults, this peptide (K**N**LSCVVK) is from the FYVE-type domain-containing protein A0A016UL41. This peptide was glycosylated in the larval sample as well, but with 8 glycoforms (Figure 4E, inset). The depicted glycan structures are based on the chitobiose core of N-glycans. The placement of fucose (red triangle) on the innermost GlcNAc is based on literature reports of core fucosylation in nematode glycans^27^, as well as the detection of similar structures by glycomics (Figure 1). The detection of di-fucosylated glycans on this peptide was unique to the adult stage, however such structures were detected on other larval proteins (Supplemental Tables 11, 12). Heterogeneity in the glycan structures decorating this peptide between stages further supports the hypothesis that glycosylation acts in addition to protein expression to modulate protein functions and interactions. The peptide with the greatest number of glycoforms in the larval sample was MQ**N**VTVK, belonging to the transthyretin-like family protein A0A0D6LL27. This protein was unique to the larval stage, but proteins from this family have been identified in the excretory/secretory proteome of *A. ceylanicum*^60^ and the related veterinary hookworm *Ancylostoma caninum*^26^. This protein also shares 32.8% identity (E value: 2e-13, query coverage: 92%) with the *H. contortus* transthyretin domain-containing protein *Hc*TTR (gi|510856026, A0A7I4YT89), which can bind IL-4^61,62^.

### Phosphorylcholine (PC)-modified N-glycans are a stage-independent feature of the *A. ceylanicum* glycoproteome

An immunomodulatory role for phosphorylcholine has been previously demonstrated in helminth infections^5–7,34,63–65^. This non-mammalian glycan modification was first detected in our glycomic analysis of adult hookworms (Supplemental Table 1). Using the intact glycoproteomic workflow, we localized PC-containing glycans to 28 larval and adult proteins (Figure 5A). A total of 11 PC-glycans decorated 38 larval glycopeptides and 28 adult glycopeptides (Figure 5B). Six of the PC-glycans were found only in one stage (2 in larvae, 4 in adults), suggesting that PC-glycan remodeling is a dynamic process that continues throughout parasite development. The identification of stage-specific non-mammalian glycans may have downstream applications as markers of infection. Of the 28 identified PC-glycoproteins, 6 were found in both stages, 12 in larvae only, and 10 in adults alone (Figure 5A, Table 2). This observation of stage specificity also aligns with our analysis of all identified glycoproteins (Figure 3C). The most common PC-glycan structures by glycopeptide count were the same in each stage: HexNAc(3)Hex(3)PC(1) (Figure 5C,D pale green) and HexNAc(3)Hex(3)dHex(1)PC(1) (Figure 5C,D pale orange), both paucimannose N-glycans. Manual validation of the raw data additionally confirmed the presence of PC. A representative glycopeptide (N**N**VTLSIDK) of an α-2-macroglobulin bait region domain-containing protein (A0A016SAS0) is shown in Figure 5C. The ability to sequence such peptides and reliably identify labile glycan modifications additionally confirms the sensitivity of our glycoproteomic workflow. To determine whether PC-glycans were a feature of a specific protein family, we performed a GO term analysis in g:Profiler (https://biit.cs.ut.ee/gprofiler/gost, Benjamini-Hochberg FDR < 0.05), however this was not the case (Supplemental Figure 2).

**Figure 5:**
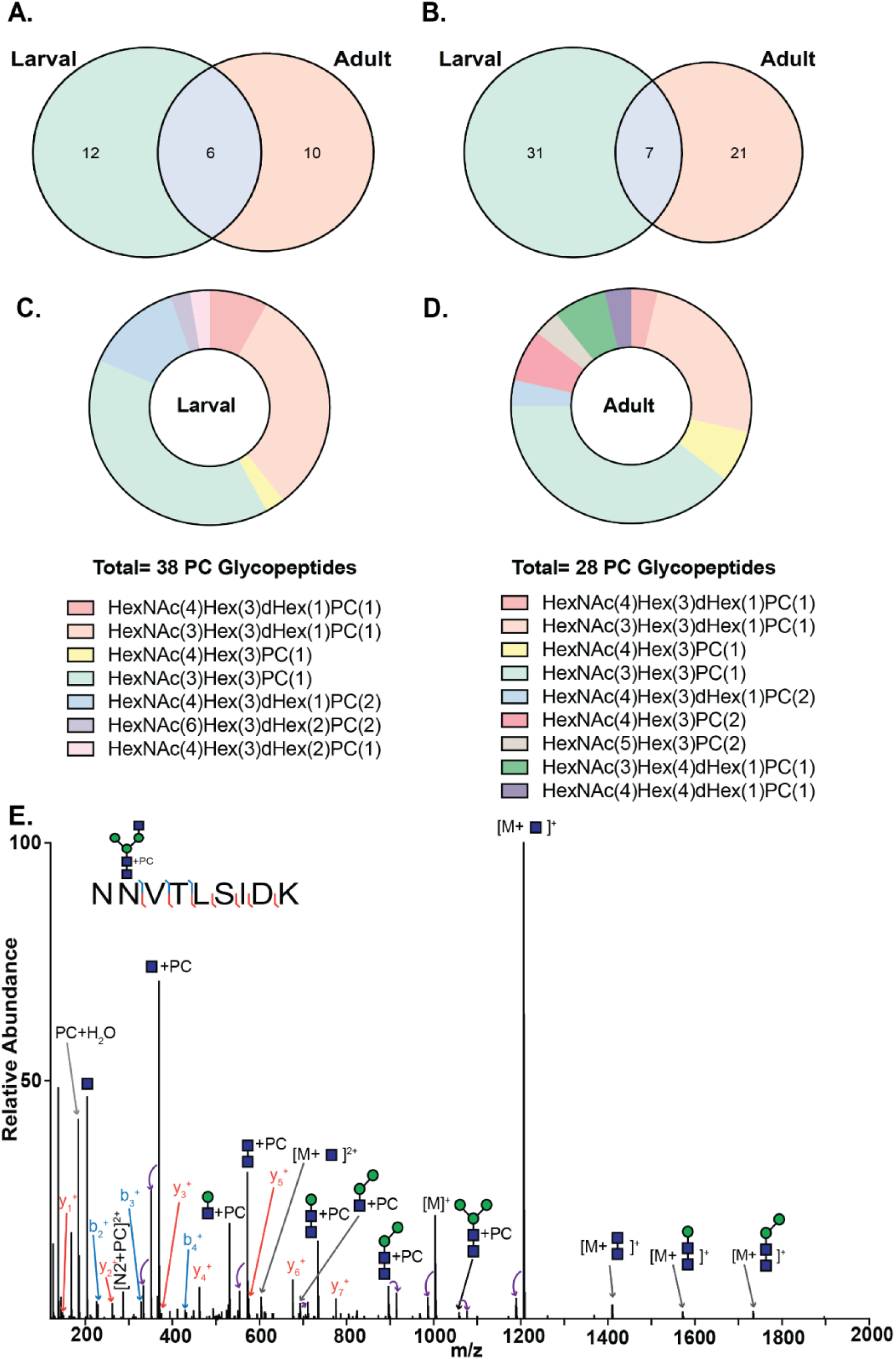
Phosphorylcholine is detected on larval and adult N-glycopeptides. Phosphorylcholine (PC) is a non-mammalian glycan modification with demonstrated immunomodulatory roles in other systems. Glycopeptides and glycoproteins with PC were identified in the final datasets for *A. ceylanicum* larvae and adults. A comparison of the PC-glycoproteins (a) and glycopeptides (b) suggest similar trends of stage specificity observed in the total set of glycoproteins (Figure 3 C,D). The distributions of glycopeptides by PC-glycan structure are shown in (c) for larval samples and (d) for adult samples. Several stage-specific structures were detected. Manual validation of a representative PC-glycopeptide (**N**VVTLSIDK) is shown in (e). Diagnostic ions for the loss of PC + H_2_O (m/z 184.0737) and PC + GlcNAc (m/z 369.1418) are annotated on the MS/MS spectrum and confirm the Byonic algorithm assignment of this structure.

**Table 2:** Hookworm N-glycoproteins identified with phosphorylcholine-containing glycans.

| Uniprot ID | Description | Parasite Stage |
| --- | --- | --- |
| A0A016SAS0 | Alpha-2-macroglobulin bait region domain-containing protein | Both |
| A0A016UL41 | FYVE-type domain-containing protein | Both |
| A0A016VA64 | Laminin EGF-like protein | Both |
| A0A016VQ95 | Transthyretin-like family protein | Both |
| A0A016W3Q3 | SXP/RAL-2 family protein Anis 5-like cation-binding domain-containing protein | Both |
| A0A016WAU9 | Fibronectin type III domain protein | Both |
| A0A016SD79 | Transthyretin-like family protein | Larval |
| A0A016SHK7 | ZP domain-containing protein | Larval |
| A0A016TJW1 | Laminin EGF-like protein | Larval |
| A0A016TVX8 | Laminin EGF-like protein | Larval |
| A0A016U1Q8 | Galectin | Larval |
| A0A016UJ94 | EGF-like domain protein | Larval |
| A0A016V6I0 | Saposin B-type domain-containing protein | Larval |
| A0A016VNC1 | BPTI/Kunitz inhibitor domain-containing protein | Larval |
| A0A016VTL5 | Uncharacterized protein | Larval |
| A0A016WRV9 | Uncharacterized protein | Larval |
| A0A016WVF4 | Fibronectin type III domain protein | Larval |
| A0A0D6LL27 | Transthyretin-like family protein | Larval |
| A0A016S4A3 | Organic solvent tolerance-like N-terminal domain-containing protein | Adult |
| A0A016S6H1 | Immunoglobulin I-set domain protein | Adult |
| A0A016S9I1 | Peptidase M13 N-terminal domain-containing protein | Adult |
| A0A016SMI1 | Tryptophan synthase beta chain-like PALP domain-containing protein | Adult |
| A0A016SYS2 | Serpin domain-containing protein | Adult |
| A0A016T1T5 | Peptidase M13 N-terminal domain-containing protein | Adult |
| A0A016TWK7 | C-type lectin domain-containing protein | Adult |
| A0A016X2C9 | Prolyl endopeptidase | Adult |
| A0A0D6LBL9 | Transthyretin-like family protein | Adult |
| A0A0D6LR74 | Serine proteinase inhibitor | Adult |

## CONCLUSION

Hookworm proteins have been studied for decades as sources of drug targets and vaccine antigens^12^. Despite a well-described role for helminth glycoproteins and glycan structures in mediating aspects of disease pathogenesis and parasite development in other systems^5,6,8,28–35,66^, virtually nothing is known about the glycosylation patterns of human hookworm proteins. To this end, we report the results of the first intact glycoproteomic analysis of the infectious (larval) and chronic (adult) stages of the hookworm *A. ceylanicum*. An initial glycomic analysis of pooled adult hookworm homogenate detected galactosylated fucose and phosphorylcholine (PC)-containing N-glycans. These structures are not characteristic of the mammalian hosts in which these obligate parasites reside, which may indicate immunologically relevant roles throughout the course of infection. Galactosylated fucose has been previously found on nematode glycans^28,47,67–70^ and can bind galectins, including those expressed in the gut^68^. In turn, PC-modified glycans are recognized by host antibodies during *Brugia malayi*^4^ and *Trichinella spiralis*^9^ infection, although PC itself may not be the dominant epitope^4^. A role for PC in immunomodulation has also been well-described^5–7,34,63–65^.

With the goal of localizing these and other N-glycans to hookworm proteins, we applied an intact glycoproteomic workflow to pooled larval and adult samples. The identified N-glycoproteins were largely stage-specific and spanned diverse molecular functions. Three glycoproteins have been previously reported in *A. ceylanicum*: neutrophil inhibitory factor^71^, and 2 putative cysteine proteases (Acey_s0154.g3007, and Acey_s0034.g2829),^72^ although their associated glycans have not been characterized. These proteins were not identified in our analyses. This may be due to excretion/secretion^71^, low expression levels, or the presence of unglycosylated glycoforms^72^ that may be favored under native conditions. Our analyses revealed that heterogeneity, in both glycosite occupancy and in the N-glycans decorating a single N_X_S/T sequon, was a dominant feature within and between stages. Such variation in N-glycosylation patterns may have consequences for host immune recognition of hookworm glycoproteins. Stage-specific N-glycans were also detected, which may have utility as markers of early or chronic infection. Additionally, in both the glycomic and glycoproteomic workflows, paucimannose glycans were frequently detected. This finding is consistent with similar reports of several nematodes, including soil-transmitted helminths^28,50,73–76^. Paucimannose glycans were recently identified on peptides of the *H. contortus* H11 vaccine antigen that bound host IgG^11^ and thus may contribute to a protective immune response. The detection of truncated, methylated, and PC-modified N-glycans also underscores the need to systematically assess recombinant expression systems for hookworm proteins. Such studies would ensure that native hookworm glycosylation patterns are reliably reproduced, and that the biological functions of these glycoproteins are accurately recapitulated.

The identification of N-glycoproteins in *A. ceylanicum* adults and larvae offers novel insight into hookworm biology and potential mechanisms of immunopathogenesis. The observed stage specificity of glycopeptides and glycoproteins further underscores fundamental differences in environmental and host niches occupied by larval and adult hookworms. These findings also suggest that the N-glycoproteome of *A. ceylanicum* is more stage-divergent than the proteome itself. As such, hookworm glycoproteins may have utility towards the development of stage-specific drugs, diagnostics, and vaccines. Further investigation of glycan-mediated interactions between *A. ceylanicum* hookworms and their hosts will be necessary to advance our understanding of this system and develop effective interventions to combat hookworm disease.

## FUNDING SOURCES

This research was made possible by CTSA Grant Number UL1 TR001863 and KL2 TR001862 or TL1 TR001864 from the National Center for Advancing Translational Science (NCATS), a component of the National Institutes of Health (NIH). Additional support was provided by the National Institute of Allergy and Infectious Diseases of the NIH through grants R01AI162826 and F31AI197571. N-Glycan profiling was performed at the Complex Carbohydrate Research Center and was supported in part by the NIH-funded R24 grant R24GM137782. The contents are solely the responsibility of the authors and do not necessarily represent the official view of NIH.

## ACKNOWLEDGEMENTS

The authors wish to acknowledge the contributions of the Yale Animal Care and Resource Center management and staff to the animal husbandry necessary for the maintenance of the *A. ceylanicum* life cycle.

## Notes

### Competing Interest Statement

The authors have declared no competing interest.

