## Supplemental Table 1, Supplemental Figures for "N-glycoproteomic profiling of the hookworm *Ancylostoma ceylanicum* reveals heterogeneity and phosphorylcholine-modified glycans in larval and adult parasites"

**Supplemental Table 1**: N-glycans identified in adult hookworm extract (HEX) by released glycan analyses. N-glycans were identified by permethylation, procainamide-labeling, or both. Proposed structures are based on MS2 data from procainamide-labeling or, for those glycans identified only through permethylation, are based on known biosynthetic pathways.

| Procainamide-labeled N-glycans $[M+Proc]^+$ | Permethylated N-glycans $[M+Na]^+$ | Composition | Proposed structure |
|---|---|---|---|
| 968.4542 | 967.444 | Hex(2)HexNAc(2) | 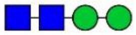 |
| 1114.5121 | 1141.510 | Hex(2)HexNAc(2)dHex(1) | 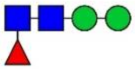 |
| 1130.5071 | 1171.517 | Hex(3)HexNAc(2) | 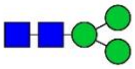 |
| 1276.5650 | 1345.573 | Hex(3)HexNAc(2)dHex(1) | 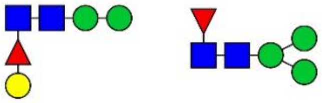 |
| 1317.5915 | 1386.589 | Hex(2)HexNAc(3)dHex(1) | 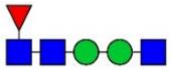 |
| 1333.5864 | 1416.595 | Hex(3)HexNAc(3) | 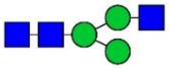 |
| 1438.6178 | 1549.636 | Hex(4)HexNAc(2)dHex(1) | 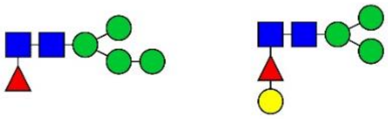 |
| 1454.6127 | 1579.780 | Hex(5)HexNAc(2) | 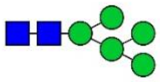 |
| 1479.6443 | 1590.658 | Hex(3)HexNAc(3)dHex(1) | 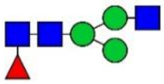 |
| 1536.6658 | 1661.680 | Hex(3)HexNAc(4) | 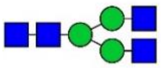 |
| 1616.6655 | 1783.704 | Hex(6)HexNAc(2) | 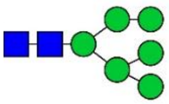 |
| 1641.6972 | 1794.898 | Hex(4)HexNAc(3)dHex(1) | 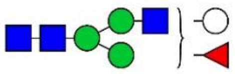 |

| | | | |
|---|---|---|---|
| 1682.7237 | 1835.749 | Hex(3)HexNAc(4)dHex(1) | 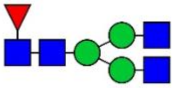 |
| 1778.7184 | 1987.982 | Hex(7)HexNAc(2) | 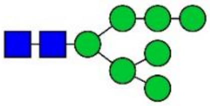 |
| 1844.7765 | 2039.827 | Hex(4)HexNAc(4)dHex(1) | 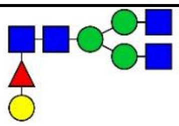 |
| 1885.8031 | 2080.854 | Hex(3)HexNAc(5)dHex(1) | 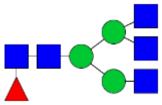 |
| 1940.7712 | 2191.871 | Hex(8)HexNAc(2) | 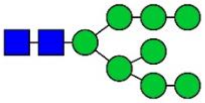 |
| 2047.8559 | 2284.933 | Hex(4)HexNAc(5)dHex(1) | 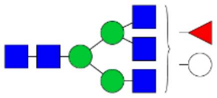 |
| 2102.8240 | 2395.969 | Hex(9)HexNAc(2) | 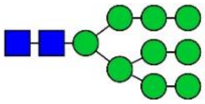 |
| - | 1519.740 | Hex(3)HexNAc(2)dHex(2) | 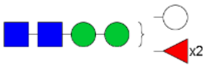  |
| - | 1906.962 | Hex(3)HexNAc(5) | 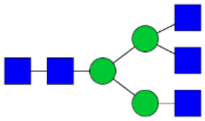 |
| - | 2243.917 | Hex(5)HexNAc(4)dHex(1) | 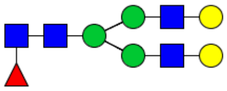 |
| - | 2605.151 | Hex(5)HexNAc(4)NeuAc(1)dHex(1) | 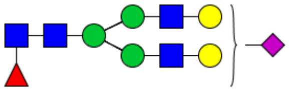 |
| - | 2723.161 | Hex(7)HexNAc(5) | 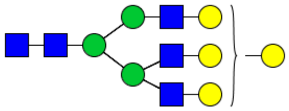 |
| 1498.6419 | - | Hex(3)HexNAc(3) + 1PC | 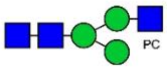  |

| | | | |
|---|---|---|---|
| 1644.6998 | - | Hex(3)HexNAc(3)dHex1 + 1PC | 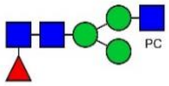 |
| 1806.7526 | - | Hex(4)HexNAc(3)dHex(1) + 1PC | 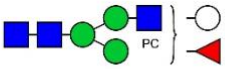 |
| 1847.7792 | - | Hex(3)HexNAc4dHex(1) + 1PC | 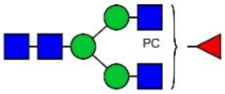 |
| 1866.7768 | - | Hex(3)HexNAc(4) + 2PC | 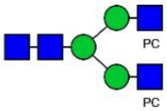 |
| 2012.8347 | - | Hex(3)HexNAc(4)dHex(1) + 2PC | 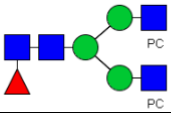 |
| 2174.8875 | - | Hex(4)HexNAc(4)dHex(1) + 2PC | 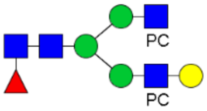 |

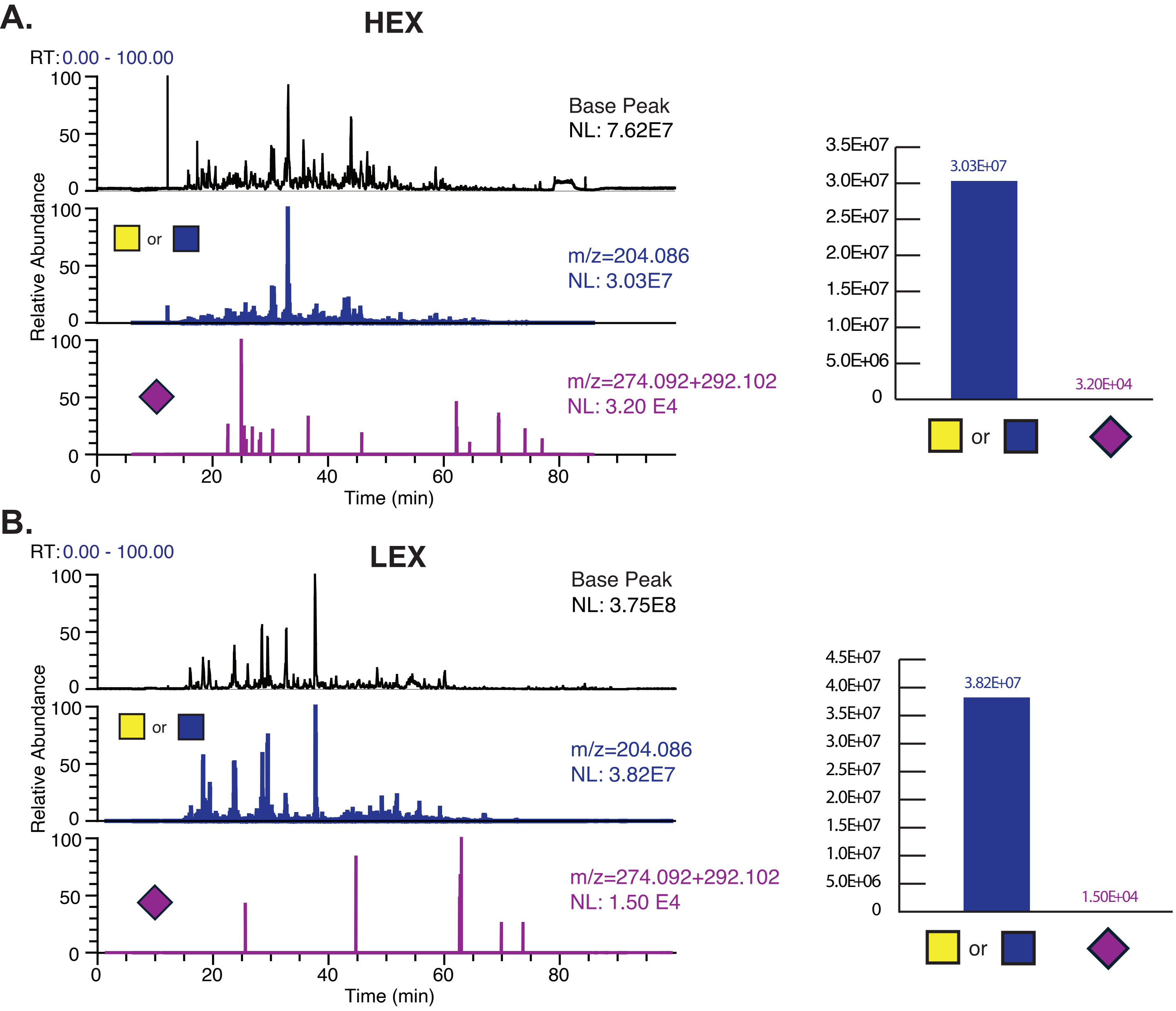


**Supplemental Figure 1:** The total base peak and oxonium ion chromatograms were extracted for a representative HEX and LEX sample. NL denotes normalized intensity level and m/z values of HexNAc and NeuAc oxonium ions are shown in blue (m/z 204.087) and purple (m/z 274.092 and 292.103) respectively. The bar graph representation of the highest intensity oxonium ion peaks are shown on the right.

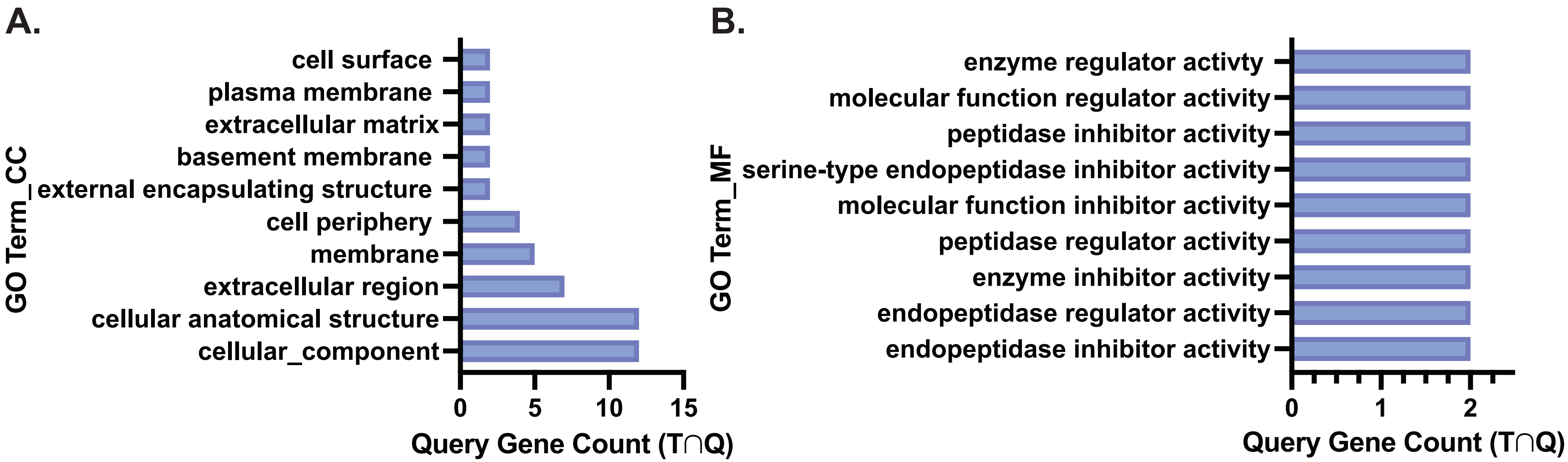


**Supplemental Figure 2**. GO Term analysis of all phosphorylcholine-containing N-glycoproteins identified from *A. ceylanicum* adults and larvae.The 28 identified glycoproteins were searched in g:Profiler against a background of all proteins identified through this workflow. Search parameters included a Benjamini-Hochberg FDR cutoff < 0.05. Results by cellular component (a) and molecular function (b) are shown above and suggest no enrichment for specific protein functions.
